# Experimental sexual selection reveals rapid evolutionary divergence in sex-specific transcriptomes and their interactions following mating

**DOI:** 10.1101/2021.01.29.428831

**Authors:** Paris Veltsos, Damiano Porcelli, Yongxiang Fang, Andrew R. Cossins, Michael G. Ritchie, Rhonda R. Snook

**Author notes:** corresponding author’s.

## Abstract

Postcopulatory interactions between the sexes in internally fertilizing species elicits both sexual conflict and sexual selection. Macroevolutionary and comparative studies have linked these processes to rapid transcriptomic evolution in sex-specific tissues and substantial transcriptomic postmating responses in females, patterns of which are altered when mating between reproductively isolated species. Here we test multiple predictions arising from sexual selection and conflict theory about the evolution of sex-specific and tissue-specific gene expression and the postmating response at the microevolutionary level. Following over 150 generations of experimental evolution under either reduced (enforced monogamy) or elevated (polyandry) sexual selection in *Drosophila pseudoobscura*, we found a substantial effect of sexual selection treatment on transcriptomic divergence in virgin male and female reproductive tissues (testes, male accessory glands, the female reproductive tract and ovaries). Sexual selection treatment also had a dominant effect on the postmating response, particularly in the female reproductive tract – the main arena for sexual conflict - compared to ovaries. This affect was asymmetric with monandry females typically showing more postmating responses than polyandry females, with enriched gene functions varying across treatments. The evolutionary history of the male partner had a larger effect on the postmating response of monandry females, but females from both sexual selection treatments showed unique patterns of gene expression and gene function when mating with males from the alternate treatment. Our microevolutionary results mostly confirm comparative macroevolutionary predictions on the role of sexual selection on transcriptomic divergence and altered gene regulation arising from divergent coevolutionary trajectories between sexual selection treatments.

## Introduction

Sexual reproduction involves both pre- and post-mating interactions between the sexes, and sexual selection influences male and female traits that mediate the fitness outcome of these interactions. While aspects of reproduction can be cooperative, the sexes can diverge over the optima of reproductive traits, such as courtship signals, fertilization and offspring production (Arnqvist & Rowe 2005). The intensity of sexual selection is linked to the extent to which reproductive fitness optima differ between the sexes and can generate sexual antagonism, in which selection acts in opposing directions on the sexes (Rice 1996; Holland & Rice 1999). As the sexes share the majority of their genome, intra-locus sexual conflict can occur over phenotypes that are influenced by these shared genes. Resolving this conflict can occur by the evolution of sexual dimorphism, such as sex-specific regulation of gene expression (Mank 2017), wherein sexual conflict is a result of outcomes over intersexual reproductive interactions from loci with sex-limited effects (inter-locus conflict). Comparative genomic studies have found that genes with rapid divergence and that show stronger signatures of positive divergent selection are often sex-biased or sex-limited in expression (e.g. Pröschel *et al.* 2006; Ellegren & Parsch 2007; Zhang *et al.* 2007; Cheng & Kirkpatrick 2016). This pattern is sometimes stronger in species with traits associated indicating strong sexual selection, such as increased sexual dimorphism (Harrison *et al.* 2015; Wright *et al.* 2019).

In internally fertilizing species, the main arena for post-ejaculatory molecular interactions is the female reproductive tract (FRT), which includes sites of sperm transfer, storage and subsequent fertilization of eggs transiting from the ovaries. Postmating female responses are extensive, influencing female behaviour, morphology and physiology. These responses are mediated by interactions between components of the male ejaculate, including sperm produced in the testes and seminal fluid proteins (Sfps) produced by accessory glands, and female reproductive proteins in the FRT and ovaries (Wolfner 2009; Sirot *et al.* 2015; Oku *et al.* 2019). Postmating changes in females include altering investment in oogenesis (Wolfner 2009), remating propensity (Chapman *et al.* 2003), and sperm storage and usage (Avila *et al.* 2010; Avila *et al.* 2015). Other aspects of female physiology are also altered, for example hunger (Carvalho *et al.* 2006), aggression (Bath *et al.* 2017) and homeostasis (Ribeiro & Dickson 2010; Cognigni *et al.* 2011). Genes associated with immunity and stress change gene expression upon mating in females, which may impact susceptibility or resistance to pathogens and/or parasites (Zhong *et al.* 2013; Oku *et al.* 2019).

An increasing number of studies have characterized transcriptomic postmating changes in females and the associated gene functions. Many studies typically examine either whole bodies or abdomens of females (Lawniczak & Begun 2004; McGraw *et al.* 2008; Innocenti & Morrow 2009; Hollis *et al.* 2014; Innocenti *et al.* 2014; Hollis *et al.* 2016; Delbare *et al.* 2017; Veltsos *et al.* 2017; Fowler *et al.* 2019). Alternatively, some studies examine only one component of the FRT, either the “lower” reproductive tract, defined by the female sperm storage organs (Mack *et al.* 2006; Prokupek *et al.* 2008) or the “upper” reproductive tract, defined by the oviducts (Kapelnikov *et al.* 2008; but see McDonough-Goldstein *et al.* 2021). Receipt of the male ejaculate effects sperm storage dynamics, oogenesis and oviposition (Sirot *et al.* 2015) making them all subject to sexual selection (and sexual conflict). Likewise, for males, the main focus on the role of sexual selection and sexual conflict has been on Sfp evolution given that they are among the most rapidly evolving proteins known (Ellegren & Parsch 2007; Ahmed-Braimah *et al.* 2017). However, sperm and the cellular architecture of the testes also can be subject to rapid morphological evolution and sexual selection (Lüpold *et al.* 2009).

The strong postmating sexual selection and sexual conflict associated with the reproductive interactions between the sexes also cause rapid evolutionary changes between lineages, which could influence reproductive isolation (Markow 1997; Manier *et al.* 2013; Ahmed-Braimah *et al.* 2020). In particular, postmating prezygotic (PMPZ) reproductive isolation in which gametes do not interact properly prior to fertilization (Ahmed-Braimah *et al.* 2017; Garlovsky *et al.* 2020) or affect egg production (Matute & Coyne 2010) is hypothesized to result from divergent coevolutionary trajectories of sexual selection and sexual conflict in isolated populations. If different populations experience different population coevolution over time, then there will be gene expression or proteomic mismatches in sexual interactions between independently evolved lineages when the male ejaculate interacts with non-coevolved female reproductive tissues (Ahmed-Braimah *et al.* 2020; McCullough *et al.* 2020; Diaz *et al.* 2021). However, such studies have focused on comparisons of the mating response of either heterospecific crosses or conspecific crosses where the sexual selection history of the populations are unknown.

The role of sexual selection in altering the coevolutionary dynamics between the sexes can be addressed experimentally. Experimental sexual selection manipulates the opportunity and strength of sexual selection by subjecting isolated populations to either polyandrous conditions, which promotes strong sexual selection, or enforced monandrous conditions, which reduces it. This approach has been used to examine the evolution of gene expression in response to sexual selection, and link it to macroevolutionary patterns of sex-biased and sex-limited gene expression evolution (Hollis *et al.* 2014; Immonen *et al.* 2014; Veltsos *et al.* 2017).

These previous studies supported the role of sexual selection in divergent sex-biased gene expression, but whether male- or female-biased genes responded the most varies between species, sexes, tissues and sexual experience, for unknown reasons (Hollis *et al.* 2014; Immonen *et al.* 2014; Veltsos *et al.* 2017; Hollis *et al.* 2019; Parker *et al.* 2019). Additionally, these studies were limited in that postmating responses and/or sex-specific tissue responses were rarely examined. Consequently, understanding how sexual selection impacts sex-limited and reproductive tissue-specific gene expression, the consequences of this divergence on postmating responses, and whether such divergence results in altered regulation of gene expression when mating between sexual selection treatments is limited.

In this study, we use replicate populations of *D. pseudoobscura* after 150 generations of experimental evolution in which either monandry is enforced (referred to as M), which reduces sexual selection and conflict or the opportunity of polyandry is elevated (referred to as E), which may increase the strength of selection and conflict to test hypotheses of the role of sexual selection on tissue-specific and sex-specific gene expression. Our previous work found substantial phenotypic responses to the manipulation of the sexual selection environment (Crudgington *et al.* 2005; Snook *et al.* 2005; Crudgington *et al.* 2009; Crudgington *et al.* 2010; Debelle *et al.* 2014; Debelle *et al.* 2016), including those that could influence postmating responses such investment of male accessory glands (Crudgington *et al.* 2009) and ovariole number and subsequent offspring production in females (Crudgington *et al.* 2010; Immonen *et al.* 2014). Sex-biased gene expression evolution has also occurred but our previous studies were based on either whole bodies of only one sex (Immonen *et al.* 2014) or heads and abdomens of each sex in the virgin and courted condition, but not following mating (Veltsos *et al.* 2017). These phenotypic responses may result from evolution of tissue- and sex-specific gene expression.

We take a quantitative transcriptomics approach to investigate the impact of sexual selection on gene expression divergence, sampling male testes and accessory glands separately, and separating the FRT into the lower reproductive tract (including the uterus and sperm storage organs) and the ovaries. We first determine whether sexual selection treatment impacts virgin gene expression in all four tissues, testing the prediction that polyandry selects for upregulation. It has previously been suggested that males subjected to intense postcopulatory sexual conflict should upregulate seminal fluid proteins for manipulation of female reproductive investment (Hollis *et al.* 2016). Likewise, polyandrous females should be poised for mating in anticipation of receipt of a manipulative male ejaculate that interacts within the FRT and thus should exhibit anticipatory upregulation of reproductive genes (Heifetz & Wolfner 2004; McGraw *et al.* 2004; Hollis *et al.* 2016). We then determine the relative impact of sexual selection, mating per se, and their interaction for each female reproductive tissue. We test the prediction that postmating interactions will diverge between sexual selection treatments, with poised polyandry females showing less upregulation upon mating relative to monandry females (see previous predictions) and that such priming will be reflected in the types of genes that alter expression, including genes acting later during oogenesis (Immonen *et al.* 2014; Veltsos *et al.* 2017) and immune and stress response genes. Immune and stress response genes have been commonly identified in female postmating transcriptomic responses, are assumed to indicate costs of mating, receipt of a foreign ejaculate and sexual conflict (Innocenti & Morrow 2009; Zhong *et al.* 2013). Mating involves interactions between the sexes so we ask whether the postmating expression response of females arises from an interaction between the sexes or is primarily driven by one sex. Given that sexual selection is stronger on males (Winkler *et al.* 2021), we test the prediction that polyandrous males will drive the female postmating transcriptomic response. Related, we test the prediction that the divergent coevolutionary trajectories exhibited between isolated monandry and polyandry populations will generate unique or more pronounced responses which could form the basis of postmating prezygotic reproductive isolation. We also test whether any effects of male sexual selection history on the female postmating response are asymmetric with the expectation that monandry females may exhibit more gene expression changes when mating with polyandry males than in the reciprocal cross. Given different coevolutionary histories of the monandry and polyandry populations, we also test the prediction that most genes induced only in non-coevolved mating will be unique to each female sexual selection treatment.

## Material and Methods

### Experimental evolution lines

The origin, establishment, and maintenance of the selection lines are described in detail elsewhere (Crudgington *et al.* 2005). Briefly, 50 wild-caught females of *D. pseudoobscura* from a population in Tucson, AZ USA were brought into the laboratory and reared for three generations, then four replicate lines of two different sexual selection treatments were established. We modified the opportunity for sexual selection by manipulating the adult sex ratio in food vials (2.5 x 80 mm) by either confining one female with a single male (enforced monogamy treatment; M, monadry) or one female with six males (elevated polyandry treatment; E, polyandry). This species is naturally polyandrous with wild-caught females frequently being inseminated by at least two males at any given time (Anderson 1974). We successfully equalized effective population sizes between the treatments (Snook *et al.* 2009). At each generation, offspring were collected and pooled together within each replicate line for each treatment, and a sample from this pool was used to start the next non-overlapping generation in the appropriate sex ratios. Thus, this proportionally reflected the differential offspring production across families within a replicate and treatment. Generation time was 28 days and all populations were kept at 22oC on a 12L;12D cycle, with standard food media and added live yeast. Note that ‘monandry’ versus ‘polyandry’ as used here refers to the evolutionary history under which the individuals have evolved, not their current reproductive status.

### Sample preparation

To generate experimental males and females, parents were collected from each replicate at generation 157-158. We standardized for maternal and larval environments as previously described (Crudgington *et al.* 2010). Briefly, parents were mated en masse in food bottles, transferred to containers with oviposition plates, allowed to oviposit for 24 h, and then 48 hr later, 100 first instar larvae were seeded in standard food vials (Crudgington *et al.* 2010). Virgin males and females were collected under light CO_2_ anaesthesia on the day of eclosion and kept separate in vials of 10 individuals for 5 days to ensure reproductive maturity (Snook 2001). On Day 5, within a 2 h window after lights turned on, one virgin female was placed in a food vial with one virgin male that was from either the same experimental replicate (“coevolved”; MM, EE where the first letter is the female) or the other treatment (“non-coevolved”; ME, EM). Our previous studies analysed gene expression in either whole body females or heads and abdomens of males and females. A potential criticism of this approach is that observed responses potentially confound changes in gene expression with allometric changes in relevant tissues (Montgomery & Mank 2016). Most importantly, we know that male and female reproductive tissues are key to evolutionary responses to sexual selection and involved in sexual interactions. Therefore, here we carried out analyses of dissected male testes, accessory glands, and female ovaries and reproductive tracts (see File S1 which illustrates the experimental design).

We dissected age- and circadian rhythm-matched virgin males and females from the same collections. Each treatment was represented by 100 individuals, the tissues of which were equally split into 4 separate tubes, for easy pooling. Each pool contained the dissected tissues of the 4 biological replicates of the E or M treatments. For the mating treatments, males were put first in individual vials with fly food and allowed to settle. Females were then added, and were dissected 6 h after the first couple mated, in the order of mating, within a 2 h block. Dissections were performed under ether anaesthesia in RNAlater (Ambion) on ice blocks. We separately collected the ovaries and the remainder of the FRT, including the sperm storage organs (seminal receptacle and spermathecae). We refer to these different female tissue sets as ovaries and the FRT. The male accessory glands and testes were also dissected separately (ejaculatory bulbs were not included). All tissues were left at 4oC in RNAlater (Ambion) for one day and then transferred to −80oC until RNA extraction. The pools were processed for RNA extraction using Trizol (Ambion) following the manufacturer’s instructions. RNA extractions were cleaned up in Qiagen RNeasy kit columns according to the manufacturer’s protocol, including the 15 min DNase treatment. The quality of RNA extractions was checked with Nanodrop and Bioanalyser.

### Sequencing and mapping

Illumina libraries were prepared using the ScriptSeq kit (Illumina Inc) following the manufacturers protocol. rRNA was depleted using Epicenters’ Ribozero kit. Paired end second stranded libraries were sequenced at 100 bp read length using an Illumina HiSeq 2000. Reads were mapped to the *D. pseudoobscura* genome v3.1, and indexed using bowtie2 (Langmead *et al.* 2009). Paired-end reads were aligned using option “-g 1 –library-type fr-secondstrand” with TopHat2.0.8b (which calls bowtie2.1.0; Kim *et al.* 2013) and instructs TopHat2 to report the best alignment to the reference. Exon features were counted using HTSeq-count (Anders *et al.* 2015) and the reads of all exons of each gene were combined to provide overall measures of gene expression (Veltsos et al. 2021).

### Statistical analyses

We analysed the count data using edgeR v3.18.1 (Robinson *et al.* 2010) running in R v.3.4.0 (R Development Core Team. 2007) with scripts from (Veltsos 2021). We combined all libraries for each tissue even when analysing contrasts that did not use all libraries. This approach allows to incorporate as much information on gene expression variance as possible, counteracting the fact that data points available for each gene in a given contrast are limited. Libraries were normalised with the TMM procedure in edgeR and only genes with at least 0 counts per million after normalization across all libraries used in each analysis, were retained. Dispersion was measured with default parameters using a negative binomial model. We considered genes to be differentially expressed (DE) if they were below the 5% false discovery rate (FDR) threshold (Benjamini & Hochberg 1995). We did not employ a log_2_FC threshold because allometry is unlikely to influence results obtained from specific tissues (Montgomery & Mank 2016).

We asked four different questions about the evolution of gene expression in response to variation in strength of sexual selection. First, we assessed whether the strength of sexual selection impacts the evolution of gene expression in virgin male reproductive tissues, by detecting differential expression in the contrast E vs M (File S2 a,b). Second, we addressed the same question in a similar manner for female reproductive tissues (File S2 c,d). Third, we investigated the effect of sexual selection history on the female mating response by analysing the contrasts equivalent to the main effect of sexual selection history (E+EE vs M+MM, where single letters indicate virgin status and double letters indicate mated status with the female partner written first), the main effect of mating (E+M vs EE+MM), and their interaction (E+MM vs M+EE). We report these results by considering the virgin gene expression status to be the baseline when compared to mated females. To separate the effects of the male and female sexual selection history on the female mating response we ran a fourth model in which we contrasted the main effect of female treatment (EE+EM vs MM+ME), the main effect of male treatment (EE+ME vs MM+EM) and their interaction (EE+MM vs EM+ME). For this analysis we consider the coevolved mating as a baseline and categorize the non-coevolved postmating response as having either relatively higher (EM or ME up) or lower (EM down or ME down) expression.

We tested the proportions of up and downregulated genes in each contrast for departure from a 50% expectation using the chisq.test function in R. When comparing the number of DE genes across virgin contrasts (Figure 1,2) which had different total numbers of genes in each contrast, we performed chisq.test on their proportion across all genes expressed in both contrasts. Finally, when comparing numbers of DE genes in the coevolved and non-coevolved treatment subsets separately for E and M mated females (Figure 4), we tested against a 50% expectation since the total number of possible genes is the same in all contrasts. This is because the contrasts are part of a model that included all 6 types of libraries for E and M females of virgin, coevolved and non-coevolved mating status. Differences in gene expression magnitude of genes within a contrast were texted using the wilcox.test function in R.

**Figure 1.**
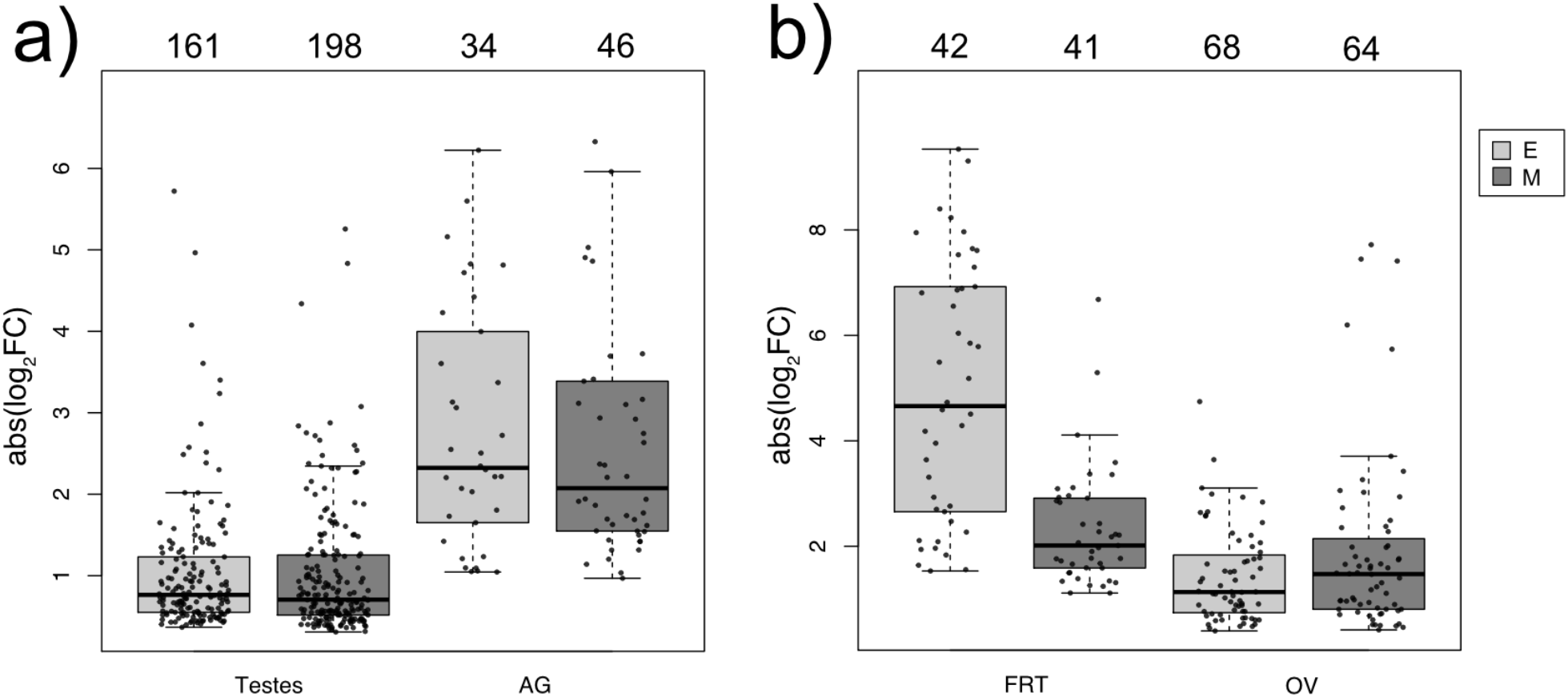
Differential gene expression (absolute log_2_FC changes, y axis) in virgin a) male or b) female tissues comparing responses of polyandry (E; light gray) or monandry (M; dark grey) selection treatments. Dots indicate DE genes and their number is noted above each box.

**Figure 2.**
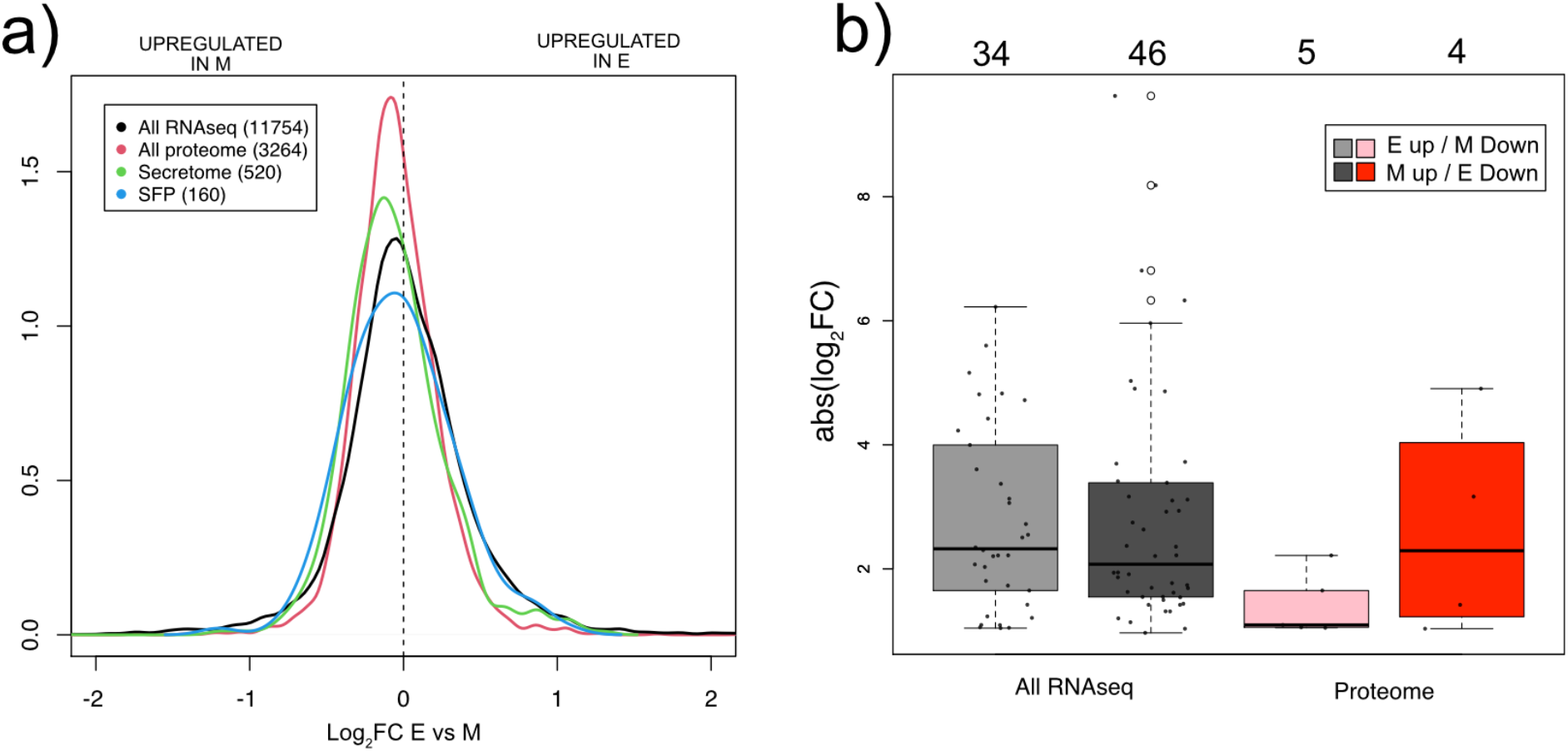
Effect of sexual selection on expression of genes coding for a) all and b) differentially expressed male accessory gland genes between polyandry (positive x axis values) and monandry (negative x-axis values) treatments. Line colors in a) differentiate between all accessory gland genes (black) or those categorized as proteome (red), secretome (green) or seminal fluid proteins (blue) based on Karr et al. 2019. The number of DE genes is indicated in the parentheses and above the boxes. Bar colors (b) differentiate the number (dots) and magnitude of absolute gene expression changes (y axis) for all DE male accessory gland genes (All RNAseq; grey) or for the accessory gland proteome genes (Proteome; colored) showing either elevated expression in E (light grey or pink) or M (dark grey or red) males.

We performed GO enrichment analysis using topGO v2.22.0 with the weight01 algorithm to account for GO topology (Alexa & Rahnenfuhrer 2010). Results with p < 0.05 on Fisher’s exact tests, corrected for topology, were retained (File S3).

For analysis of Sfps, we contrasted the distribution of the change in expression (log_2_FC) of all genes, and DE genes in the contrast between sexual selection treatments for virgin accessory gland transcriptomes using density plots and boxplots, respectively. We compared the distributions of three Sfp-related gene subsets, identified from *D. pseudoobscura* proteomics (Karr *et al.* 2019). The largest subset was 3281 proteins produced in the accessory gland (“proteome”). Of these, 528 had protein secretory signals (“secretome”) and 163 were also orthologous to *D. melanogaster* seminal fluid proteins (putative Sfps or “exoproteome”). The majority of these genes could be cross-referenced to our data (Figure 2 legend), but only proteome genes were detected among the DE accessory gland genes.

## Results

### Sexual selection causes gene expression divergence in virgin male reproductive tissues

Previous work has hypothesized that monandry selects for downregulation and polyandry selects for upregulation of genes expressed in male reproductive tissues in the abdomen of *D. melanogaster* (Hollis *et al.* 2016). We found that sexual selection history caused divergence in gene expression in virgin male testes for 359 DE genes. However, contrary to prediction, there were marginally more upregulated genes in monandry (Figure 1a; x^2^=3,81, df=1, p<0.051; File S2). Those 198 genes were related to the biological processes (BP) of proteolysis, digestion and the innate immune response (File S3; tab “testis EvsM”). Τhe 161 upregulated genes in polyandry males had BP enrichment related to stress responses, such as double strand break repair, cellular response to UV, and terpenoid metabolism (File S3; tab “testis EvsM”).

In male accessory glands, 80 genes were DE between sexual selection treatments but there was no difference in the number of upregulated genes between treatments (Figure 1a; x^2^=1,8, df=1, p<0.18). The 34 accessory gland genes upregulated in polyandry males were enriched for BP terms related to eggshell chorion assembly, development, and neuropeptide signaling (File S3; tab “agland EvsM”) whereas the 46 genes upregulated in monandry males were enriched for the BP term “detection of chemical stimulus involved in sensory perception” (File S3; tab “agland EvsM”).

Despite Sfps being among the most rapidly evolving proteins, putatively in response to sexual selection, we found a larger proportion of testes genes was differentially expressed compared to accessory gland genes (359/11751 vs 80/11754; x2=179.47, df=1, p<0.001). The magnitude of gene expression within each tissue did not differ between monandry and polyandry males (testis: W = 14712, p = 0.21; accessory glands: W=752, p = 0.78; Figure 1a). To examine whether Sfps had higher expression in polyandrous males, as predicted (Hollis *et al.* 2016), we compared expression levels between the sexual selection treatments of different sets of accessory gland proteins recently described for *D. pseudoobscura* (Karr *et al.* 2019). We did not support the prediction; monandry males had higher overall expression of secretome proteins compared to polyandry males (W=21345000, p<0.001; Figure 2a) and sexual selection treatment did not affect gene expression of proteome or Sfp genes. Examination of the 80 accessory gland genes that were differentially expressed between treatments, regardless of set, we found a non-significant trend towards higher expression in polyandry males (Figure 2b).

In conclusion, virgin male reproductive tissue expression was affected by experimental manipulation of the strength of sexual selection but not in the predicted direction. Both the total number and proportion of DE genes is higher in testes. There also was a weak trend for monandry, not polyandry, males to upregulate testes genes and accessory glands with secretory signals.

### Sexual selection causes gene expression divergence in virgin female reproductive tissues

Similar to males, we tested whether monandry selects for downregulation and polyandry selects for upregulation of genes (Hollis *et al.* 2016) in virgin female reproductive tissues (FRT and ovaries), such that polyandry females are more poised for subsequent mating. While there was an effect of sexual selection on gene expression in both tissues, the number of genes upregulated in each sexual selection treatment did not differ (Figure 1b; FRT: x^2^=0,01, df=1, p =0,9; ovaries: x^2^=0,12, df=1, p =0,73). In the FRT, 42 genes were upregulated in polyandry and were enriched in several BP terms associated with the immune system whereas 41 genes upregulated in monandry were enriched for insemination (File S3; tab “FRT EvsM”). In ovaries, the 68 genes upregulated in polyandry, but not the 64 genes upregulated in monandry, were enriched in BP terms associated with eggshell chorion assembly (File S3, tab “ovEvsM”).

The proportion of DE genes in ovaries was significantly greater than in the FRT (132/8625 vs 83/10273; x^2^=21.12, df=1, p<0.001). Even though there was no difference in the number of upregulated genes between sexual selection treatments, the magnitude of gene expression was impacted in a tissue-specific way. In the FRT, but not ovaries, logFC of upregulated genes in polyandry was greater than in monandry (FRT: W = 302, p <0.001; ovaries: W = 2452, p = 0.21; Figure 1b).

Overall, we find that experimental sexual selection alters gene expression and, while there is no treatment bias in number of responding DE genes, FRTs have a higher magnitude of upregulation in polyandry compared to monandry. Moreover, the reproductive tissues respond differently, with a higher percentage of ovarian genes changing in expression compared to the FRT. These results support the hypothesis that polyandry females are more poised for mating via increases in expression magnitude in the FRT, the primary site of molecular interactions between the female and male ejaculate.

### Sexual selection causes divergence in the female postmating response

With regards to the female postmating response, we expect polyandry females to already express postmating response genes as virgins, whereas monandry females would increase the expression of these genes after mating (the poised hypothesis; Heifetz & Wolfner 2004; McGraw *et al.* 2004). This response would be seen as a significant interaction in our model (Table 1).

**Table 1.**
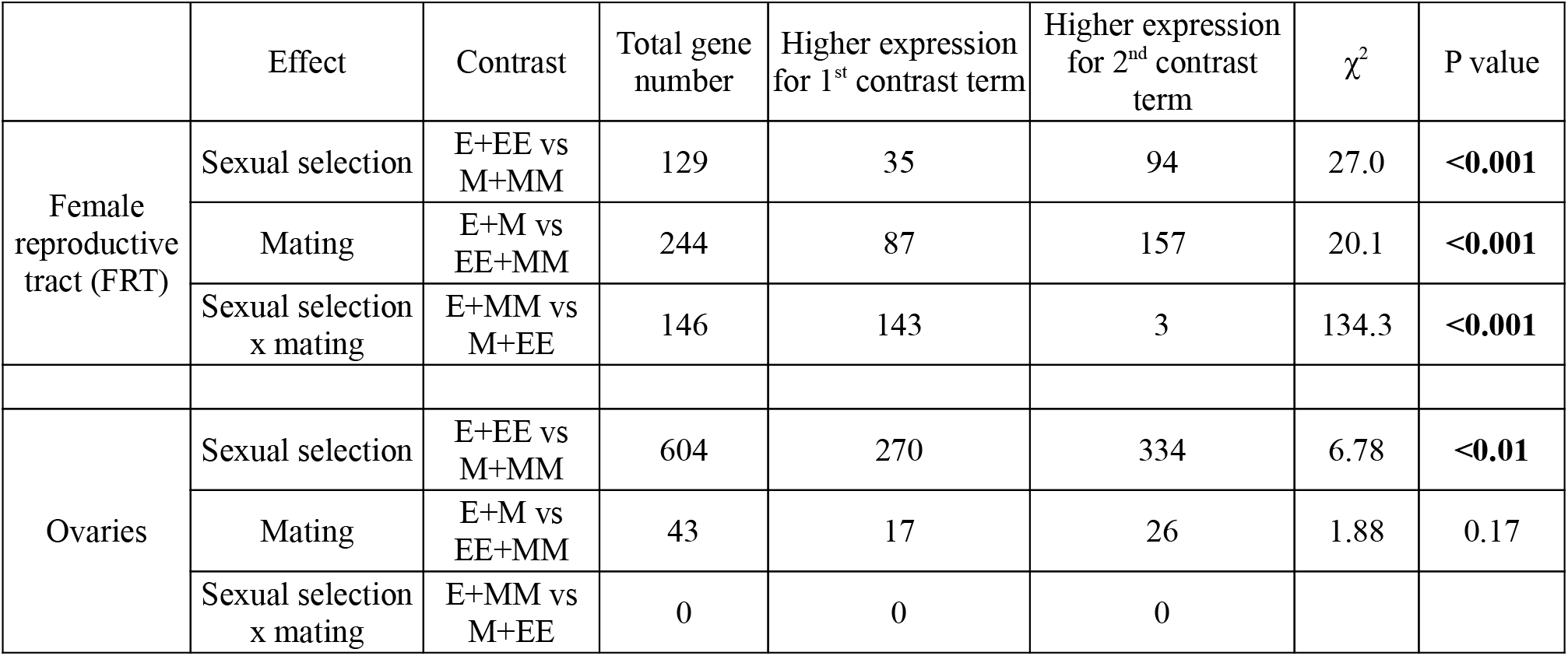
Outcome of contrasts testing the effect of sexual selection, mating status, and their interaction on the female postmating response in the female reproductive tract (FRT) and ovaries. Df are always 1. Chi-square tests were performed against the null hypothesis of 50% for the number of upregulated genes between the first and second contrast terms. Statistically significant results are indicated in bold. Single-letter contrast names indicate virgins, two letters indicate mated, with the female partner written first.

For the FRT, we find both main effects of sexual selection treatment and mating, and their interaction, to be significant (Table 1). To illustrate the interaction, we plot the main effect of mating within each female treatment on separate axes, and separately indicate genes that always respond to mating and those that show significant interaction effects between mating and sexual selection treatment (Figure 3a). Most (143/146) of the significant genes for the interaction were upregulated upon mating in monandry females (and downregulated upon mating in polyandry females; Table 1; Figure 3a [bottom right quadrant]). These genes were enriched for immune function, a variety of metabolic processes, and eggshell chorion assembly (Figure 3b; File S3; tab “Interact treatment x mating FRT”). The 3 genes showing upregulation in mated polyandry and downregulation in monandry have BP terms related to reproduction (Figure 3b; File S3; tab “Interact treatment x mating FRT”). Of the 244 DE genes responding to mating regardless of sexual selection history, more were upregulated after mating than downregulated (Table 1, Figure 3a). The commonly upregulated genes after mating were enriched for BP terms associated with stress and immune responses, only one of which was shared with BP enrichment of genes significant for the interaction (Figure 3b; File S3; tab “Main effect of mating FRT”). Genes downregulated after mating were not enriched for BP related to immune/stress responses (Figure 3b; File S3; tab “Main effect of mating FRT”). With regard to the main effect of sexual selection treatment (Table 1), 94 genes were upregulated in monandry (but showed no BP enrichment) and only 35 genes were upregulated in polyandry (with two enriched BP terms - defense response to fungus and ecdysteriod metabolic process; File S3; tab “Main effect of SS line FRT”; Figure 3b).

**Figure 3.**
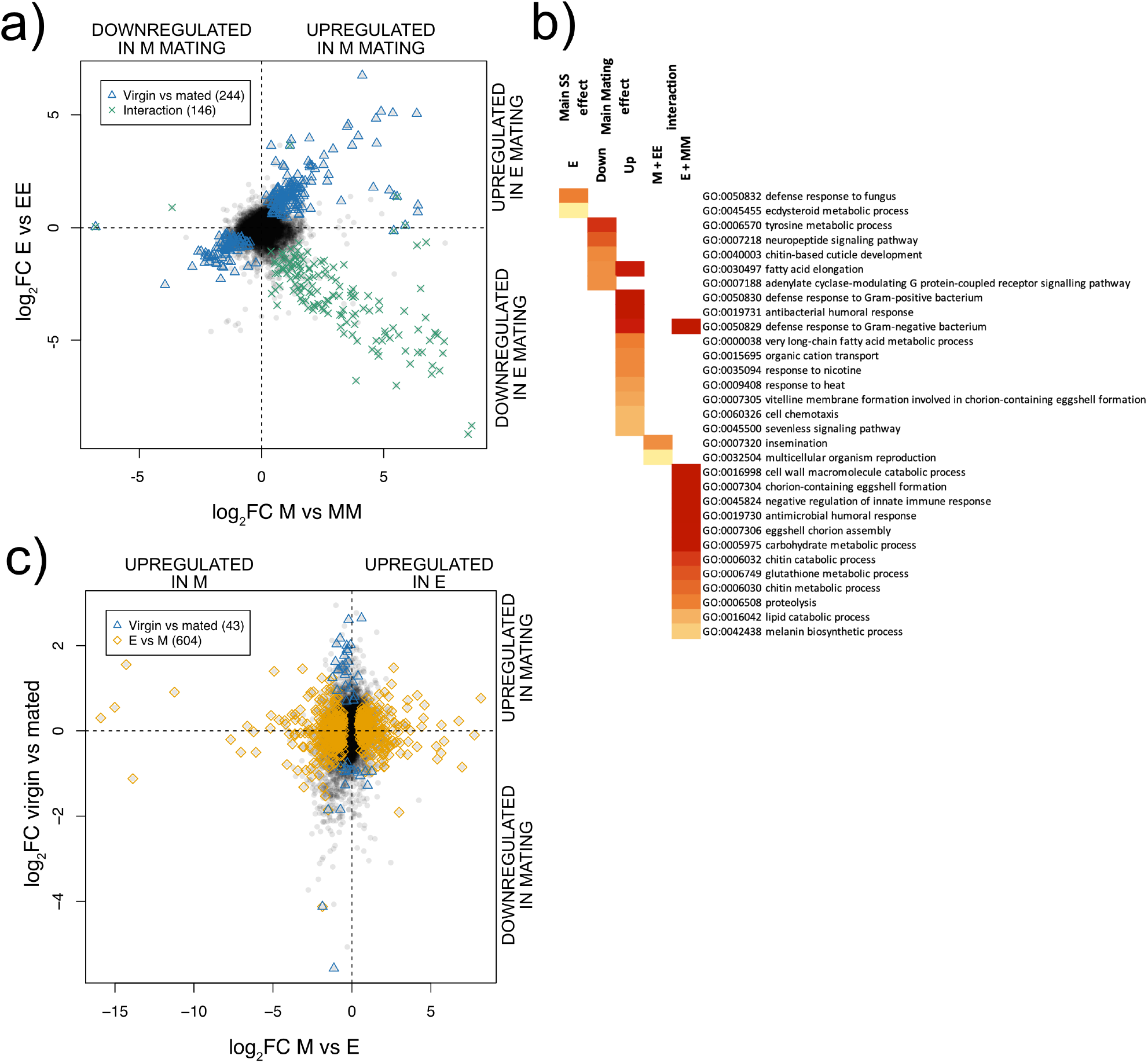
Postmating gene expression (log_2_FC) responses and enriched GO BP terms in female reproductive tissues. A) Significantly DE genes in female reproductive tracts (FRT) with DE genes indicated with blue triangles for the main effect of mating status, yellow diamonds for the main effect of sexual selection treatment and green crosses for the interaction between the two. The axes show the coevolved postmating response for each sexual selection treatment, which allows to visualize the interaction between sexual selection treatment and mating as the diagonal on which the green crosses fall on. B) Significant BP term enrichment for the main effect and interactions in FRT. C) the coevolved postmating response in ovaries with axes representing the two main effects, as there were no significant genes for the interaction. The number of DE genes is indicated in the parentheses. See Table 1 for statistical basis.

In the ovaries, only the main effect of sexual selection treatment was associated with differential gene expression. As with the FRT, there were more upregulated genes in monadry vs polyandry (Table 1; Figure 3c). The 334 genes upregulated in monandry females were enriched in BP terms related to tissue development whereas the 270 genes upregulated in polyandry were enriched for eggshell chorion assembly (File S3; tab “Main effect of SS treatment OV”).

Thus, analyses for both the FRT and the ovaries suggest that polyandry females are more poised for mating based on reduced number of DE genes after mating. This interpretation is further supported by BP enrichment of upregulated genes under polyandry that relate to egg production in ovaries and stress and immune responses in the FRT. The latter may attest to polyandry males being potentially immunogenic (Innocenti & Morrow 2009).

### Female, not male, sexual selection treatment drives the female postmating gene expression response

The postmating gene expression response in female reproductive tissues represents an interaction between the sexes. To examine these interactions in more detail, we partitioned gene expression in females mated to males of the same or different sexual selection treatment, determining the effect of female treatment, male treatment, and their interaction. The interaction tests whether matings between coevolved individuals with respect to sexual selection treatment respond differently than matings between non-coevolved individuals (Table 2). Since the strength of sexual selection is stronger on males than females (Winkler *et al.* 2021), we predicted that both the male effect and the interaction would have the strongest impact on the female postmating response (Table 2).

**Table 2.** Outcome of contrasts testing the effect of female sexual selection treatment, male sexual selection treatment, and their interaction on the female postmating response in the female reproductive tract (FRT) and ovaries. Df are always 1. Chi-square tests were performed against the null hypothesis of 50% for the number of upregulated genes between the first and second contrast terms. Statistically significant results are indicated in bold. NA indicates the numbers were too low to meaningfully statistically compare. Single-letter contrast names indicate virgins, two letters indicate mated, with the female partner written first.

| | Effect | Contrast | Total gene number | Higher expression for 1<sup>st</sup> contrast term | Higher expression for 2<sup>nd</sup> contrast term | $\chi^2$ | P value |
| --- | --- | --- | --- | --- | --- | --- | --- |
| Female reproductive tract (FRT) | Female | EE+EM vs MM+ME | 198 | 38 | 160 | 75.2 | **<0.001** |
|  | Male | EE+ME vs MM+EM | 7 | 0 | 7 | NA | NA |
|  | Female x Male | EE+MM vs EM+ME | 0 | 0 | 0 | NA | NA |
| Ovaries | Female | EE+EM vs MM+ME | 566 | 306 | 206 | 19.5 | **< 0.001** |
|  | Male | EE+ME vs MM+EM | 0 | 0 | 0 | NA | NA |
|  | Female x Male | EE+MM vs EM+ME | 6 | 4 | 2 | NA | NA |

Contrary to predictions, in both female tissues, we found no significant interaction effect and little to no male effect (Table 2). There was a small male effect in the FRT with 7 DE genes, enriched for egg chorion BP, all upregulated after mating with monandry males (Table 2; File S3; tab “Main male effect FRT”). Surprisingly, for both the FRT and ovaries, the female effect was stronger and the sexual selection effect asymmetric between tissues. Of the 198 DE genes in FRT, 160 were upregulated when monandry females mated (Table 2) and are enriched for immune response and metabolic processes, while the genes upregulated in polyandry females did not show any significant enrichment (File S3; tab “Main female effect FRT”). In contrast, in the ovaries, of the 566 DE genes, 306 were upregulated in mated polyandry females and these genes were enriched for eggshell chorion assembly BP. The 206 genes upregulated after mating in monandry females did not have GO enrichment terms consistent for any clear function (File S3; tab “Main female effect OV”).

In summary, female sexual selection history plays a critical role in determining the extent and function of the postmating response, with the effect of male and interaction between the sexes minimal. Again, polyandry females had suppressed gene expression changes relative to monandry females in the FRT. However, in the ovaries, polyandry females had more upregulated genes although these were related to egg production, suggesting that polyandry females can “gear up” for oogenesis more quickly than monandry females. One caveat to this analysis is that it combines females from the two sexual selection treatments to test for male and female effects. Given that we showed above that sexual selection treatment influences the female postmating response, consequences of interactions between males and females may be obscured.

### Sexual selection asymmetrically alters gene expression

To further decompose the effect of sexual selection treatment origin on the female postmating response, we examined gene expression in coevolved and non-coevolved matings separately for females of different sexual selection history. This also allowed us to test the prediction that divergent coevolutionary trajectories between sexual selection treatments would generate unique or more pronounced responses when mating with a non-coevolved male. We predict this effect will be asymmetric as monogamous females have not evolved in a highly manipulative environment in response to high male competition.

In the ovaries, both polyandry and monandry females had limited postmating changes in gene expression, regardless of the sexual selection treatment of the male partner (Table 3). Combined with the insignificant overall effect on mated females (Table 2), this suggests that the males have a reduced and similar effect on the postmating response of ovaries in monandry and polyandry females. In contrast, patterns were varied in the FRT, in which monandry females showed significant gene upregulation following mating, regardless of mating partner (Table 3) whereas in polyandry females there was no significant postmating response when mating with males of either sexual selection treatment (Table 3).

**Table 3.** Outcome of contrasts testing the effect of male origin on the female postmating response in the female reproductive tract (FRT) and ovaries for monandry and polyandry females. Df are always 1. Chi-square tests were performed against the null hypothesis of 50% for the number of upregulated genes between the first and second contrast terms. Statistically significant results are indicated in bold. NA indicates the numbers were too low to meaningfully statistically compare. Single-letter contrast names indicate virgins, two letters indicate mated, with the female partner written first.

| | Effect | Contrast | Total gene number | Higher expression for 1<sup>st</sup> contrast term | Higher expression for 2<sup>nd</sup> contrast term | $\chi^2$ | P value |
| --- | --- | --- | --- | --- | --- | --- | --- |
| Female reproductive tract (FRT) | Monandry coevolved | M vs MM | 160 | 57 | 103 | 13.3 | **<0.001** |
|  | Monandry non-coevolved | M vs ME | 363 | 157 | 206 | 6.61 | **<0.01** |
|  | Polyandry coevolved | E vs EE | 199 | 99 | 100 | 0.005 | 0.94 |
|  | Polyandry non-coevolved | E vs EM | 90 | 51 | 39 | 1.6 | 0.21 |
| Ovaries | Monandry coevolved | M vs MM | 1 | 0 | 1 | NA | NA |
|  | Monandry non-coevolved | M vs ME | 3 | 1 | 2 | NA | NA |
|  | Polyandry coevolved | E vs EE | 6 | 1 | 6 | NA | NA |
|  | Polyandry non-coevolved | E vs EM | 0 | 0 | 0 | NA | NA |

Divergent coevolutionary trajectories between sexual selection treatments may result in unique gene expression between coevolved and non-coevolved crosses, irrespective of whether the total number of genes alter expression. Indeed, while we found some genes alter gene expression in the same direction regardless of male mate origin, many genes are uniquely expressed dependent on male origin and this is asymmetric across sexual selection treatments. For monandry females, more unique genes were expressed when mating with polyandry males than when mating with coevolved males (coevolved = 55 vs non-coevolved = 258; x^2^=131.66, p<0.001; Figure 4a) but the opposite pattern occurs in polyandry females (coevolved = 139 vs non-coevolved = 30; x^2^ =70.3, p<0.001; Figure 4c). Comparing the female sexual selection treatments, mating with non-coevolved males caused more genes to alter expression in monandry females than in polyandry females (monandry = 258 vs polyandry = 30; x^2^ =180.5, p<0.001; Figure 4a,c). Regardless of DE number, uniquely DE genes for non-coevolved crosses show a unique pattern of gene expression for both sexual selection treatments (Figure 4; compare the red x’s to the purple dots and blue crosses), which generally do not overlap with either shared or uniquely coevolved responses.

**Figure 4.**
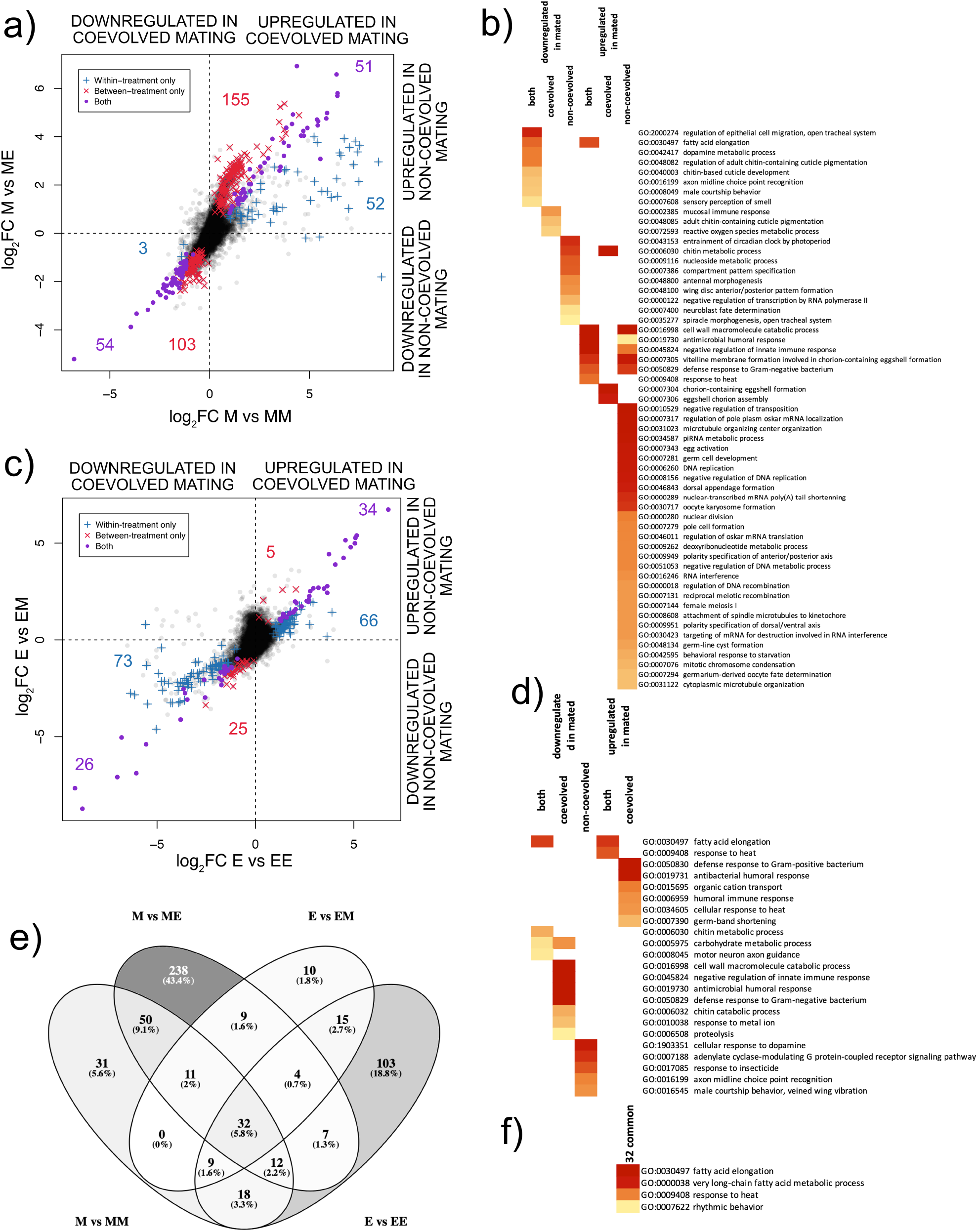
Postmating responses of DE genes (log_2_FC) in the female reproductive tracts (FRT) of a) monogamy and c) polyandry females when mated to males either from the same sexual selection treatment or the opposite treatment. Genes are categorized as being DE only in coevolved mating (blue +), only in non-coevolved mating (red x) or regardless of male treatment (purple o). In each plot, the x axis represents is the coevolved postmating response and the y axis the non-coevolved postmating response with negative values representing virgin-biased genes and positive values representing mating-biased. The colored numbers within the plots are the number of DE genes for each gene category. Panels b) and d) represent the associated GO BP enrichment of the DE genes of panels a) and c) respectively. Panel e) shows the overlap of genes corresponding to coevolved and non-coevolved mating separately in monogamy and polyandry females, and panel f) shows the GO BP enrichment of the 32 consistently DE genes all e) contrasts.

Given novel genes were altered in expression when mating with noncoevolved males, we also examined GO terms for the genes that differ in expression between coevolved and non-coevolved crosses. For monandry females, many of the BP terms for uniquely upregulated genes after non-coevolved mating are related to DNA replication and germ cell development, and downregulated non-coevolved responses are related to morphogenetic processes (Figure 4b; File S3; tab “venn FRT M”). Genes affected by the male partner in polyandry females had few BP enrichment terms with unclear biological interpretation (Figure 4b; File S3; tab “venn FRT E”). Moreover, the identity of postmating response genes is unique when comparing across monandry and polyandry females, even when mated with non-coevolved males (Figure 4e). Only 32 DE genes are shared across all four mating combinations, suggesting that these are critical to female postmating responses (Figure 4e; represented by some of the purple dots in Figure 4a,c). These shared genes are enriched for four BP processes associated with fatty acid production, stress response and rhythmic behavior (Figure 4f; (File S3; tab “venn 32”).

Overall, these results confirm predictions, highlighting consistent asymmetric female postmating responses between sexual selection treatments. Results suggest monandry females are less resistant to male manipulation and/or polyandrous males are more manipulative and/or polyandrous females are more resistant, that differentially regulated genes in coevolved and non-coevolved matings showed little overlap in enriched BP terms between females of the different sexual selection treatments, and that, despite differences in total number and function, non-coevolved responses had different expression patterns than coevolved crosses.

## Discussion

Understanding how sexual selection impacts the molecular basis of sexual interactions is important because it can have profound effects on sex-specific fitness and is predicted to influence the evolution of postmating prezygotic reproductive isolation. Here we combined transcriptomics with experimental evolution to determine how sexual selection affects gene regulation simultaneously in multiple sex-specific tissues in male and female virgins and in the postmating female response when mating with either males that evolved under the same or different sexual selection treatment. Sexual selection and sexual conflict should result in polyandry males that are manipulative and females that are resistant whereas enforced monogamy should reduce male manipulation and female resistance (Arnqvist & Rowe 2005). We test whether signatures of these predictions can be identified in changes in gene regulation over the time frame of 150 generations of experimental manipulation of sexual selection and whether short term changes in the female postmating response also diverges.

We tested the prediction that polyandry selects for upregulation in virgin male and female reproductive tissues (Heifetz & Wolfner 2004; McGraw *et al.* 2004; Hollis *et al.* 2016). Given the role of accessory glands in manipulating female reproductive investment, and that the FRT is the main site of molecular interactions between the sexes, we expected that these tissues would show more divergent gene regulation than testes or ovaries. Support for these predictions varied. In males, sexual selection resulted in divergent gene regulation in both testes and accessory glands. Contrary to predictions, a larger proportion of genes change expression in the testes and relaxation of sexual selection resulted in more upregulated genes in the testes and higher expression of accessory gland genes with a signal of a secretory function. However, the function of genes that changed expression in the testes and accessory glands were distinct for each sexual selection treatment, suggesting differences in the potential consequences on male reproductive fitness and effects on female mates. In females, sexual selection resulted in divergent gene regulation in both the FRT and ovaries, but a larger proportion of ovary genes were differentially expressed compared to the FRT. These results are contrary to predictions. However, the magnitude of differential gene expression is larger in the FRT and for polyandry females, which supports the prediction. Previously we showed that virgin polyandry *D. pseudoobscura* females have more ovarioles than monandry females (Immonen *et al.* 2014). Using female whole body microarrays, we found upregulated genes in polyandry females to be associated with oogenesis (and likely to be expressed in the ovary) whereas upregulated genes in monandry were associated with genes in somatic tissues and metabolism (Immonen *et al.* 2014). In the current analysis, virgin polyandry females had upregulated genes with functions in immune responses and later stages of egg production whereas virgin monandry females upregulated genes associated with BMP signalling pathway which is involved in early patterning the *Drosophila* eggshell (Niepielko *et al.* 2012). These results support the hypothesis that polyandry females are poised for receipt of a potentially manipulative ejaculate (Heifetz & Wolfner 2004; McGraw *et al.* 2004).

We also examined the female postmating response testing the effect of sexual selection, mating and their interaction on gene expression divergence. In particular, we tested the prediction that polyandry females would show a smaller postmating response relative to monandry females given polyandry females are poised for mating. We additionally asked whether postmating responses are specific to sexual selection treatment with respect to biological processes. As predicted, we found an interaction between sexual selection treatment gene expression and mating status but this was tissue specific, occurring only in the FRT, and asymmetric across sexual selection treatments, with most DE genes upregulated in mated monandry females. While gene expression in ovaries showed no interaction effect, monandry females also upregulated more genes after mating than polyandry females, with gene functions supporting more advanced reproductive development in polyandry females (also supported in Immonen *et al.* 2014; Veltsos *et al.* 2017). Overall, these results support the hypothesis that polyandry females are poised for mating. In the FRT, DE genes associated with the interaction effect and for mating status were enriched for immune function, a commonly observed effect of mating in Drosophila (Innocenti & Morrow 2009; Sirot *et al.* 2015; Hollis *et al.* 2019). Previously, it has been suggested that upregulation of these genes arises from sexually antagonistic interactions between the sexes, such that males are immunogenic to females (Innocenti & Morrow 2009; Zhong *et al.* 2013). However, there remains insufficient data to determine whether these effects are detrimental or beneficial to females overall (Oku *et al.* 2019; Bagchi *et al.* 2021).

Given that sexual selection changes the female postmating response in both female reproductive tissues, that the female postmating response arises as an interaction between the sexes, and that sexual selection is stronger on males (Winkler *et al.* 2021), we expected that effects of male treatment and interactions should drive the female postmating transcriptomic response. This should be particularly prominent in the FRT, as the main arena for between-sex reproductive molecular interactions. We tested these predictions by determining the relative contributions of females, males and interactions between the sexes on the female postmating response. However, disentangling male and female effects is difficult within each sexual selection treatment so, to test these predictions, we crossed males and females within and between sexual selection treatments. Contrary to predictions, we found no interactions and little male effect, in both the FRT and ovaries. The dominant or only effect was attributed to female sexual selection treatment and this effect varied between tissues. The FRT of monandry females showed significantly more upregulated genes (enriched for immune function) than for polyandry females whereas, in the ovaries, polyandry females upregulated significantly more genes (enriched for later stage egg production) than monandry females. These results are consistent with our other analyses showing that polyandry females are more poised for mating and reproduction with little effect of male partner.

This latter result is surprising and may be confounded by the test performed, which combines non-coevolved and coevolved crosses in estimating each sex effect. We independently tested the monandry and polyandry female postmating response based on male partner origin to test the hypothesis that gene expression divergence, mediated by variation in sexual selection between isolated populations, generates potential reproductive mismatches via altered gene expression regulation of female postmating responses. Moreover, given that monandry should relax male manipulation and female resistance, we tested the prediction that altered regulation when mating with a noncoevolved male would be asymmetric across the different sexual selection treatments and, because of different evolutionary trajectories of monandry and polyandry populations, target different types of genes between the sexual selection treatments. Such responses may eventually generate postmating prezygotic (PMPZ) reproductive incompatibilities. Several studies have compared postmating transcriptomic responses to test the idea of reproductive mismatches generating PMPZ (Bono *et al.* 2011; Ahmed-Braimah *et al.* 2020; McCullough *et al.* 2020). However, it is difficult to infer the historical role of different evolutionary processes from patterns of contemporary divergence between species and therefore whether mismatches generated PMPZ or evolved after divergence cannot be determined. Experimental evolution can address this problem as it can distinguish between current and historical processes, but lacks the full complexity of natural conditions and typically does not result in reproductive isolation over the relatively short time frame studied.

Using experimental sexual selection, we find patterns that support these predictions in the FRT, but not in ovaries. Gene regulation in the FRT was sensitive to male treatment and varied between monandry and polyandry females. We inferred altered regulation by the identification of uniquely DE genes when mated with a non-coevolved male and comparing their prevalence with uniquely DE genes when mated with a coevolved male. Both monandry and polyandry females had more unique DE genes when mated to polyandry males than monandry males, suggesting polyandry males can manipulate female gene expression. Monandry females exhibit more DE genes when mating with polyandry males, suggesting that monandry females are less resistant to male manipulation, as predicted. The uniquely upregulated genes in monandry females from mating with a non-coevolved male were associated with DNA replication and germ cell development suggesting polyandry males manipulate investment in reproduction. Interestingly, the fewer genes affected by the male partner in polyandry females showed no clear affected process. Finally, uniquely DE genes from non-coevolved crosses showed different expression patterns than shared or uniquely DE coevolved patterns as would be expected given no recent coevolutionary history. It remains to be determined whether continued divergence and these mismatched gene regulations would generate PMPZ.

In conclusion, our results test several predictions arising from sexual selection and sexual conflict theory, and highlight substantial gene expression divergence both in the long term following 150 generations of altered sexual selection intensity and short term plastic responses when mating. We found sex- and tissue-specific effects of sexual selection on gene expression and gene function, alterations in gene expression and gene function specific to origin of the female and male partners, and predicted asymmetric altered gene regulation and function arising from divergent coevolutionary trajectories between sexual selection treatments. Changes in gene expression we identified here and in sex-biased gene expression in response to sexual selection (Veltsos *et al.* 2017) have recently been shown to associate with genomic divergence in these lines (Wiberg *et al.* 2021). Overall then, our results complement work in natural populations in which sexual selection has been implicated in gene expression and genomic divergence.

## Acknowledgements

This work was supported by the Natural Environment Research Council (NE/I014632/1 to M.G.R., A.R.C., and R.R.S), the Natural Environment Research Council Biomolecular Analysis Facility (NBAF654 to M.G.R), and the Swedish Research Council (Vetenskapsrådet; 2018-04598 to R.R.S). We thank the many people that have contributed to the generation and maintenance of the experimental sexual selection lines in RRS’s lab, R Axel W Wiberg for help with dissections, and Chris Wheat and Rachel Steward for ideas to represent GO terms. We also thank 3 anonymous reviewers for comments that substantially improved the manuscript.

## Data Access

The RNAseq data underlying this article have been submitted to ArrayExpress under accession number E-MTAB-10047. The scripts to reproduce the analysis are available at https://osf.io/z7fm9/?view_only=054171ba3f534f839a0814fa1b8f9f61.

## Author Contributions

The idea was developed by MGR and RRS, with funds obtained by MGR, RRS and ARC. The study design was contributed to by PV, MGR, RRS, ARC and YF. Biological material was prepared by RRS, PV and DP. Data were analysed by YF and PV. The first draft of the manuscript was written by PV, MGR and RRS. The final manuscript was read, edited and approved by all authors.

## Supporting Information

Additional supporting information in may be found online in the Supporting Information section.

## Supplementary Files

**File S1:**
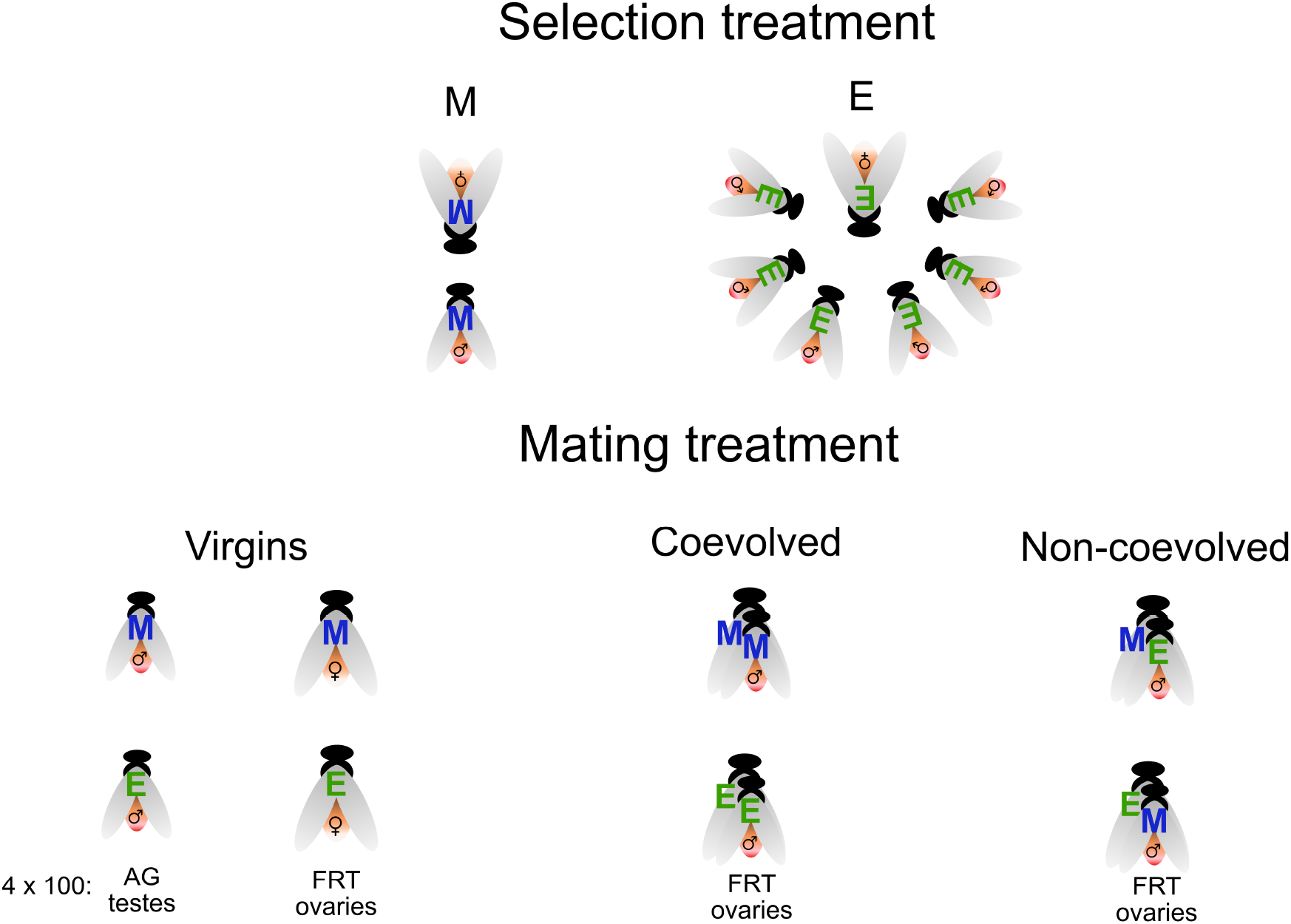
Illustration of the experimental design showing both the long-term selection treatment, and the short-term mating treatment. M: monandry, E: elevated polyandry.

**File S2:**
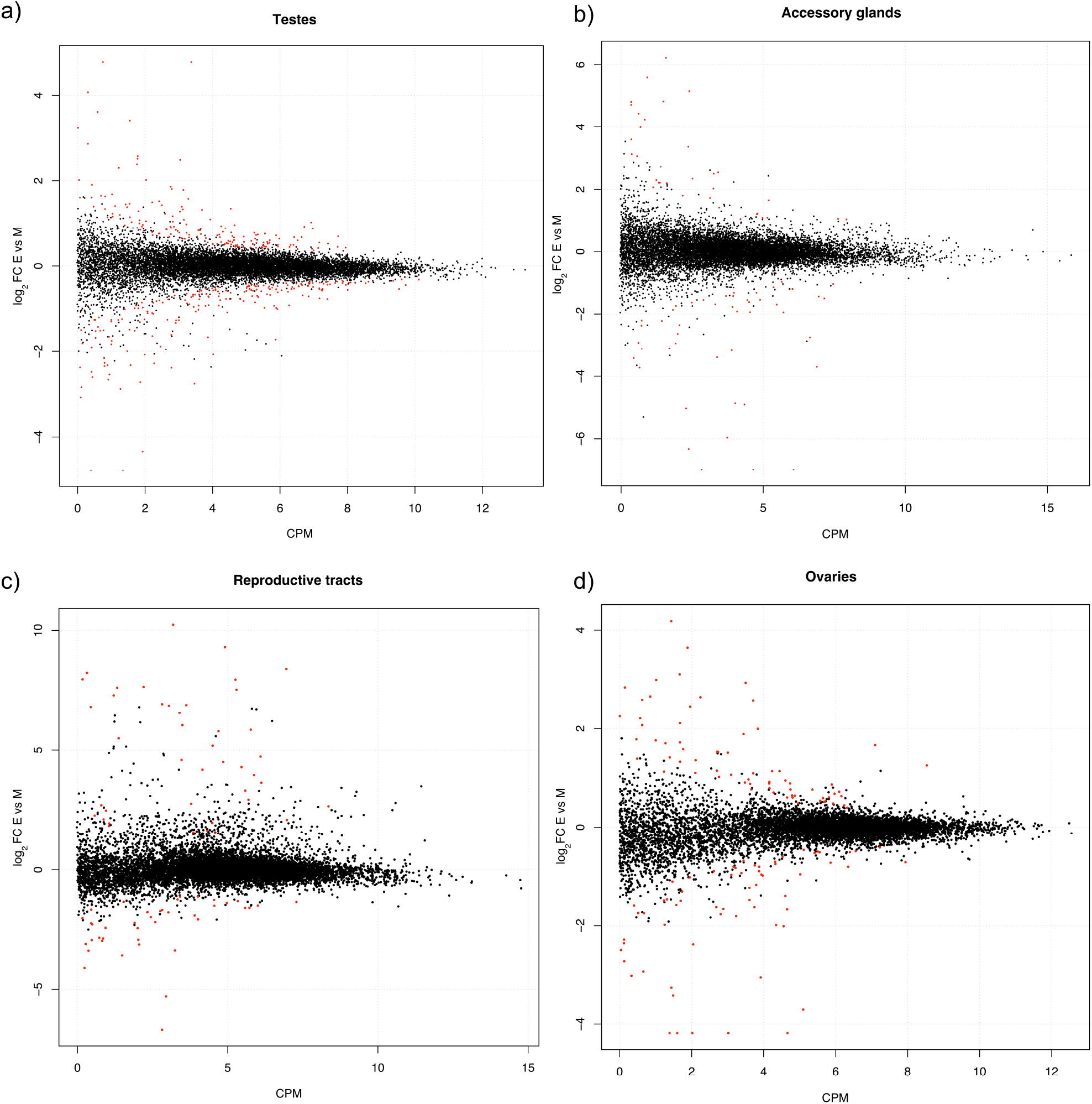
MA plot indicating DE genes in high and low (E vs M) sexual selection contrasts in all tissues for virgin flies. Genes upregulated in E have positive values on the y axis.

**File S3**: Summary of GO terms for all analysed DE genes.

